# Fluoxetine Delivery for Wound Treatment Through an Integrated Bioelectronic Device – Pharmacokinetic Parameters and Safety Profile in Swine

**DOI:** 10.1101/2025.03.11.642735

**Authors:** Anthony Gallegos, Houpu Li, Hsin-ya Yang, Guillermo Villa-Martinez, Itidal Bazzi, Samhitha Sathyanarayanan, Narges Asefifeyzabadi, Prabhat Baniya, Wan Shen Hee, Mohammadhossein Siadat, Elizabeth Chang, Sriansh Pasumarthi, Elham Aslankoohi, Mircea Teodorescu, Marcella Gomez, Marco Rolandi, Rivkah Isseroff

**Author notes:** Correspondence to: Rivkah Isseroff, UC Davis Department of Dermatology, 3301 C Street Suite 1400, Sacramento, CA 95816, USA. Correspondence to: Marco Rolandi, UC Santa Cruz Department of Electrical and Computer Engineering, 1156 High Street SOE3, Santa Cruz, CA 95064, USA. E-mail addresses (R.R. Isseroff), (M. Rolandi).

## Abstract

Wound infections are a significant medical challenge, often leading to chronicity or systemic infection. Selective serotonin reuptake inhibitors (SSRIs) have emerged as potential non-antibiotic candidates with demonstrated ability to limit growth and biofilm formation in Gram-negative bacteria, in addition to their pro-healing activity. Here, we compared direct delivery of the SSRI fluoxetine by topical bolus dosing to delivery from an iontophoresis bandage device with an actuator for temporally controlled drug delivery, in a porcine excisional wound model. Device delivery of fluoxetine resulted in a maximum concentration of 12.25 ng fluoxetine per mg tissue, compared to 2.926 ng/mg following bolus dosing, and tissue fluoxetine levels were higher after application using the device than after bolus dosing across the range of doses tested (p=0.0041). The half-life of fluoxetine in the wound tissue was 0.988 ± 0.256 days. Fluoxetine was not detected in the pig plasma, and plasma serotonin levels were not affected by the topical application. Fluoxetine delivery using the device, but not bolus delivery, produced tissue concentrations above the minimum inhibitory concentration (MIC) for some clinically important species of bacteria.

The experimental device can effectively deliver topical fluoxetine to the wound, producing higher tissue concentrations of fluoxetine at lower cumulative doses compared to bolus dosing, and with minimal risk of off-target effects. The device may simplify wound treatment by reducing the burden for daily drug application, possibly increasing adherence to a prescribed treatment regimen.

## 1. Introduction

Skin wounds, from injury, surgical procedures, and underlying medical conditions, are a common medical condition, and wound care poses a major burden to the healthcare system in the US and worldwide[1,2]. A retrospective analysis of Medicare beneficiaries in the US for the 2014 calendar year determined that 14.5% of beneficiaries (8.2 million patients) were diagnosed with wounds, with or without infection, and that infected surgical wounds were the most common category (4.0% of beneficiaries) followed by infected diabetic wounds (3.4% of beneficiaries) and nonhealing surgical wounds (3.0% of beneficiaries). A conservative estimate of the cost of treatment for acute and chronic wounds was $31.7 billion; this estimate only reflects care received through the Medicare system, and the cost is expected to be substantially higher if patients using other payment systems are included[1].

Wound infections present a significant medical challenge, complicating healing and often leading to outcomes such as chronic wounds or systemic infections[3,4]. The rise of microbial resistance to antibiotics further exacerbates this problem, creating an urgent need for alternative strategies[5,6]. To that end, investigators have sought to expand the armamentarium of treatment options by repurposing existing drugs which have demonstrated antimicrobial activity[7], an approach that affords benefits such as faster regulatory approval and reduced costs[8]. Selective serotonin reuptake inhibitors (SSRIs), commonly used for the treatment of psychiatric disorders, have emerged as potential candidates[9–11], and recent work has established the minimum inhibitory concentrations (MICs) of the SSRI fluoxetine in combination with antibiotics against several clinically important species of bacteria[12–14].

Previously, we showed that fluoxetine reduced biofilm formation in *S. aureus* infected human ex-vivo wounds, and enhanced healing in *S. aureus* infected mouse excisional wounds[15]. In addition to potential antimicrobial effects, fluoxetine has been shown to have pro-reparative effects, improving skin wound healing[15–19]. Studies have demonstrated enhanced keratinocyte migration and neovascularization, reduced expression of inflammatory cytokines, and a shift in macrophage polarization towards a less inflammatory and more pro-reparative phenotype. For these reasons, we chose fluoxetine as the drug to be delivered in this study.

Recently developed endogenously responsive drug delivery systems (DDSs) are activated by signals from the wound, such as temperature[20–22] or chemical signals[23–26]. In such devices, the rate of drug release depends primarily on the characteristics of the encapsulating matrix[27]; in addition, physiological signals may not be sufficiently strong to trigger drug release at the appropriate time[28]. To remedy this, several groups have developed exogenously responsive drug delivery systems which provide greater control over the dose and timing of drug release. Systems have been designed to respond to the programmed application of electric fields[29], temperature[30], and light[31,32]. However, there is a need for devices which allow for highly precise and fully programmable control of drug delivery[28]. To this end, we developed a programmable, wirelessly controlled, wearable wound management device containing a voltage driven iontophoretic actuator for delivery of fluoxetine and an integrated high-resolution wound imaging system. The device is controlled remotely via wireless commands sent from a laptop, using a software module with a user-friendly graphical interface, and the treatment program is carried out in real-time.

The goal of the current work is to compare the pharmacokinetics of topically applied fluoxetine when delivered using the experimental device to conventional bolus pipette delivery. We hypothesize that topical delivery can produce effective concentrations of fluoxetine in the wound without systemic perturbation, and that metered delivery using the device will optimize absorption into the wound tissue. Using a porcine wound model, wound tissue concentrations were interrogated after topical bolus dosing or application using the device, and pharmacokinetic parameters of fluoxetine in the wound were established. To examine the potential for off-target effects, pig plasma was interrogated for systemic fluoxetine and norfluoxetine, and for changes in serotonin. In addition, we interrogated the wound tissues for serotonin to examine the ability of topical fluoxetine to modulate skin serotonin levels.

### 1.1 Device design and principles of operation

The experimental device contains an iontophoresis pump actuator and wireless imaging module in a wearable format and is described in detail in our other work[33]. A schematic of the device is shown in Fig. 1.

**Fig. 1.**
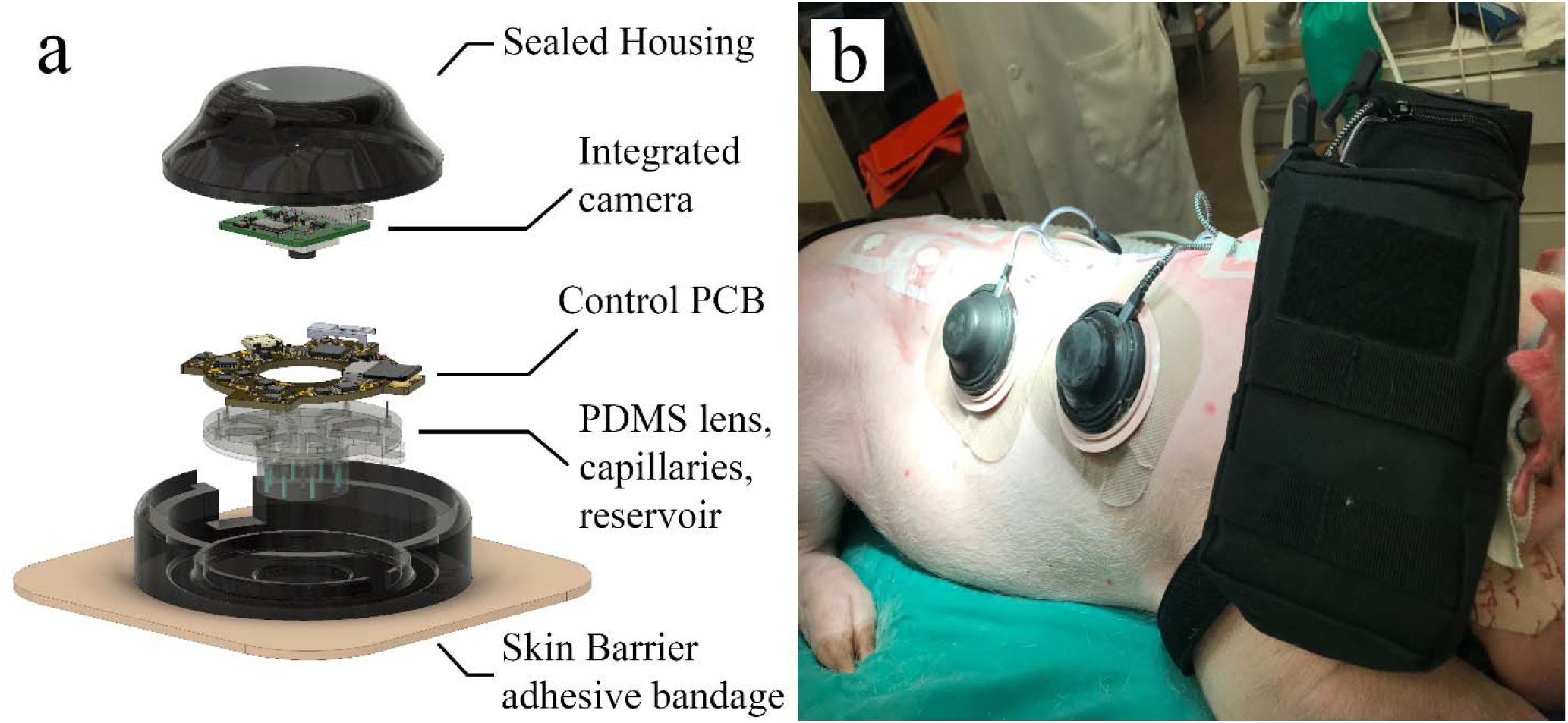
(a) An exploded diagram of the experimental device. (b) The experimental device applied to pig dorsal wounds in an experimental setting. Batteries are stored in pouches attached to the shoulder harness.

The actuator is voltage-driven, fully programmable, and allows for precise delivery of fluoxetine ions to the wound. The polydimethylsiloxane (PDMS) base contains reservoirs with microchannels for ion transport. Hydrogel ion exchange membranes are embedded within glass capillaries in the PDMS and serve as interfaces between the device and the wound, selectively delivering fluoxetine and preventing counterion transfer. Silver wires in the reservoirs connect to a custom-printed circuit board (PCB) for device control, described in our other work[34,35], and the circuit is completed by a counter electrode and the wound bed.

The PCB controls the applied voltage, V_FLX_, which drives fluoxetine ions from the reservoir into the wound via the working electrode. The PCB also manages wireless communication to an external computer for real-time monitoring and data transmission. V_FLX_ is adjustable from zero to 4.7 V through a microcontroller and digital-to-analog converter (DAC).

The imaging module is a high-resolution camera positioned at the center of the PDMS actuator, and the clear bottom of the actuator serves as a lens. The camera captures images of the wound bed which are transmitted wirelessly to a computer, allowing for continuous monitoring of the wound. The actuator and camera are enclosed in a protective casing and affixed to a commercially available adhesive bandage.

The device supports wireless operation, and commands from a computer regulate both the ion pump and imaging processes. By combining targeted drug delivery with real-time monitoring, the device is designed to enhance wound healing in clinical and experimental settings.

## 2. Materials and methods

### 2.1 Pig acclimation and wounding surgery

Yorkshire-Landrace-Duroc cross breed pigs (28-87 kg, 9-12 months old, female) were purchased from the UC Davis Swine Research Center. All animal work was approved by the UC Davis Institutional Animal Care and Use Committee (protocol #23353). Animals were acclimated for 7-10 days and trained for harness use and human contact as previously described[36]. Surgical preparation and wounding were as described[36], with the modification of prophylactic intravenous administration of Cefazolin, 30-35 mg/kg, during surgery. Baseline blood samples were collected and stored for plasma assays.

Six to ten circular wounds of 20 mm diameter and 6-10 mm depth were created on the paravertebral area of the pig as previously described[36], rinsed with 0.001% cefazolin solution, and dressed as previously described[36] for pipette and standard of care treatments, or with the experimental devices. The devices were connected to a V75 battery (Voltaic Systems, New York, USA) and Raspberry Pi 4B controller (Raspberry Pi Ltd., Cambridge, England), carried in 2 Molle pouches (Tacticool Firearms, Virginia, USA) attached to the shoulder harness of the pig. The post-operative analgesic buprenorphine, 0.005-0.01 mg/kg, was given by intramuscular injection before extubation, and a fentanyl patch was applied at the end of surgery. Pigs were monitored for signs of hyperthermia after recovery.

### 2.2 Fabrication of the wearable PDMS device

The PDMS actuator contains three main components which were molded separately: the reservoir, the notch, and the channel piece. The molds were 3D printed using a Formlabs 3B printer with Model V3 resin at a layer thickness of 50 um. To improve optical performance, an additional mold was made from CNC-milled acrylic disks with a 19.5 mm diameter.

PDMS (Sylgard 184) base and cross-linker were mixed in a 10:1 ratio. The mixture was degassed, poured into the molds, leveled, and degassed again. The molds were cured at 60°C for 48 hours. Once cured, the PDMS components were cleaned in isopropanol (IPA), rinsed with water, and oven-dried for assembly.

Silver wires were inserted into the PDMS reservoirs, and PDMS seals were applied at the insertion points to prevent leaks. The device was assembled using plasma bonding; surfaces were exposed to plasma for 10 seconds and then cured for 30 minutes at 60°C. Bond quality was tested by injecting air into the reservoirs.

Adhesive tape was applied to protect the optical window, and parylene coating was performed over the course of three hours to produce a durable protective layer. The PCB was attached to the top of the PDMS, electrodes were soldered to the corresponding pins on the PCB, and a chlorinated silver wire was attached to serve as the counter electrode. A second parylene layer was applied for additional protection.

### 2.3 Preparation of the hydrogel capillaries

Glass capillaries of 0.800 mm internal diameter were cleaned with NaOH and conditioned with deionized water, silane coupling agent, and ethanol. Once dry, the capillaries were filled with a hydrogel precursor and UV-cured at 7 mW/cm^2^. The cured hydrogel was cut into 5 mm segments and soaked in fluoxetine solution.

### 2.4 Capillary insertion and final assembly

The device and capillaries were sterilized in a biosafety cabinet using UV radiation for 30 minutes, and the remaining steps were performed in a biosafety cabinet using aseptic technique. The capillary segments were inserted into the actuator channels using a biopsy punch, and the reservoirs were filled with fluoxetine solution. The channels were sealed using 3D-printed biocompatible caps. The assembled device was mounted onto a New Image Flat CeraPlus Skin Barrier (Hollister, Illinois, USA) for attachment to the wound site, and the finished devices were placed into autoclaved steel bins for transport to the surgery room.

### 2.5 Device startup and daily operation

Upon startup, the device entered standby mode and connected to a designated Wi-Fi network in the vivarium. Researchers controlled the device remotely via wireless commands sent from a laptop, using a software module with a user-friendly graphical interface. The laptop also ran software modules for wound stage prediction[37] and closed loop control using a machine-learning algorithm[38], which were tested in a different set of pig experiments.

The device operated for 22 hours daily with a 2-hour standby window for data transfer, battery replacement, and routine checks. The battery supported up to 36 hours of operation and was replaced daily. After each battery change the device data were copied to the laptop’s hard drive, and the software module was manually reset. This was repeated daily during the experiment.

### 2.6 Fluoxetine delivery

Experimental devices were examined daily, and devices were replaced every 3-4 days during the experiments. Devices were programmed to deliver a target dose of 0.45 mg/wound/day within a six-, twelve-, or twenty-two hour treatment window. The actual dose delivered was calculated using current measurements recorded by the device controller. The actuator was programmed to stop automatically when the treatment target was reached or when the preset duration ended.

Wounds that received bolus topical fluoxetine solution, bolus saline control solution, or standard of care were examined daily, and dressings were replaced every 3-4 days or as needed. Sterile fluoxetine hydrochloride solution was prepared in saline and applied in doses of 0.45 mg/wound (‘high’ dose) or 0.025 mg/wound (‘low’ dose) on each day of application.

At the end of observation on post-operative days 3, 7, 10, or 22 (for device experiments) or on days 7 or 10 (for pipette experiments), plasma was collected, and the animal was euthanized. Wound tissue was excised with a 10 mm margin of surrounding intact skin, divided for HPLC, and flash-frozen using liquid nitrogen. Wound tissue was stored at -80°C until use.

### 2.7 Chromatographic analysis of fluoxetine and norfluoxetine

Measurements were performed using reverse phase HPLC with UV absorbance detection. The system was a Neurotransmitters Analyzer (Antec Scientific, Leyden, Netherlands) coupled to a BlueShadow 40D detector (Knauer, Berlin, Germany). Separation was performed on an Acquity UPLC BEH C18 column (3.0 mm ID x 100 mm L, 1.7 um particles, Waters, Ireland) at a flow rate of 0.450 mL per minute and column oven temperature of 35°C. The mobile phase consisted of 35:65 acetonitrile: 45 mM phosphate buffer pH = 6.0, and column effluent was monitored at 230 nm.

### 2.8 Chromatographic analysis of serotonin

For serotonin analysis in the plasma, measurements were performed using reverse phase HPLC with UV absorbance detection. The system was an Antec Neurotransmitters Analyzer coupled to a Knauer BlueShadow 40D detector. Separation was performed on a Waters Acquity UPLC BEH C18 column (3.0 mm ID x 100 mm L, 1.7 um particles) at a flow rate of 0.450 mL per minute and column oven temperature of 37°C. The mobile phase consisted of 4:96 acetonitrile: 50 mM citrate buffer pH = 4.3, and column effluent was monitored at 275 nm.

For serotonin analysis in the pig wound tissues, measurements were performed using HPLC with electrochemical detection. The system was an Antec Neurotransmitters Analyzer fitted with a 2 mm glassy carbon working electrode and Ag/AgCl reference electrode (SenCell, Antec Scientific), operated in the DC mode at a potential of 0.460 V versus Ag/AgCl. Separation was performed on a Waters Acquity UPLC BEH C18 column (1.0 mm ID x 100 mm L, 1.7 um particles) at a flow rate of 0.050 mL per minute and column oven temperature of 37°C. The mobile phase consisted of 4:96 acetonitrile: phosphate-citrate buffer pH = 6.0, containing a total of 100 mM phosphate, 100 mM citrate, 0.1 mM EDTA, and 600 mg/L octanesulfonic acid.

### 2.9 Extraction of fluoxetine and norfluoxetine from pig plasma

Pig plasma samples were purified on C18 solid-phase extraction (SPE) cartridges (100 mg, 3 mL tube volume) packed into glass tubes with glass fiber frits (Macherey-Nagel, Duren, Germany). Plasma extraction was carried out as follows: on a vacuum manifold, cartridges were conditioned with two tube volumes methanol and one tube volume water. Pig plasma (0.500 mL) was spiked with 50.0 uL internal standard solution (40.0 ug/mL fluvoxamine in methanol), diluted with 0.500 mL H_2_O, and passed through the conditioned SPE cartridges. Next, cartridges were rinsed with one tube volume H_2_O followed by one tube volume 50:50 methanol: H_2_O. Fluoxetine and norfluoxetine were eluted from the cartridges in 0.500 mL methanol containing 0.5% (v/v) formic acid, the eluate was made up to 1.000 mL by addition of 0.500 mL H_2_O, and 10 uL was injected for analysis.

### 2.10 Extraction of serotonin from pig plasma

Pig plasma samples were depleted of protein as follows: pig plasma (0.100 mL) was spiked with 5.00 uL antioxidant solution (1.3% (m/v) ascorbic acid and 2 mM EDTA in H_2_O) and 20.00 uL internal standard solution (300.0 uM N-methylserotonin in H_2_O). Next, 5.64 uL of 70% perchloric acid solution (Thermo Scientific, Waltham, Massachusetts) was added and the samples were vortexed for 30 seconds. The samples were centrifuged, and 10 uL of the supernatant was injected for analysis.

### 2.11 Tissue cryo-pulverization

Pig wound tissues were briefly thawed and trimmed to a 2 mm margin of healthy skin at the wound edges. Fat was trimmed from the bottom of the tissue pieces, and large wounds were bisected. Tissues were diced using a razor blade, were immediately refrozen, and were crushed in a stainless-steel die (BioSpec Products, Bartlesville, Oklahoma) over liquid nitrogen. Using a porcelain mortar and pestle over liquid nitrogen, the crushed tissues were ground to the consistency of a fine powder. Frozen ground tissue was stored at -80°C until analysis.

### 2.12 Extraction of fluoxetine and norfluoxetine from pig wound tissues

Ground pig wound tissues (20.0 to 30.0 mg) were resuspended in 0.250 mL methanol containing 0.5% (v/v) formic acid and 2.000 ug/mL fluvoxamine. The resuspended tissues were sonicated for 15 minutes in a sonicating bath, and the resulting tissue homogenates were centrifuged. The supernatants were transferred to clean microcentrifuge tubes and made up to 0.500 mL by addition of 0.250 mL H_2_O, and 10 uL of this was injected for analysis.

### 2.13 Extraction of serotonin from pig wound tissues

Ground pig wound tissues (20.0 to 30.0 mg) were resuspended in 10 uL of 0.1 N HClO_4_ per mg tissue and sonicated for 10 minutes in a sonicating bath. The resulting tissue homogenates were centrifuged, passed through 0.2 um nylon syringe filters (Thermo Scientific, Waltham, Massachusetts), loaded into pre-rinsed 30 kDa ultrafiltration cartridges (Pierce, Thermo Scientific) and centrifuged at 15,000 x G for 30 minutes. 3 uL of the filtrate was injected for analysis.

### 2.14 Statistical analysis

Statistical analyses were performed in GraphPad Prism 10.2 software. Linear correlations were evaluated using Spearman’s test for nonparametric data, and hypothesis testing for the comparison of regression lines was performed using ANCOVA. Test results of p ≤ 0.05 were considered statistically significant. Outliers were identified using the ROUT function in Prism 10.2 software with a false discovery rate of Q = 2%.

## 3. Results

### 3.1 Fluoxetine delivery by the experimental device increases wound tissue concentrations compared to bolus pipette delivery

Pig wound tissue was collected after 3 or 7 days of daily topical fluoxetine application using either the experimental device or a pipette and was analyzed by HPLC to determine the fluoxetine tissue concentration. For both application modalities, the tissue level of fluoxetine was strongly correlated to the cumulative dose delivered (device, Spearman’s R=0.862, p=0.0089; pipette, R=0.956, p=0.022). Norfluoxetine, the major active metabolite of fluoxetine, was not detected in any of the wound tissues (data not shown). Application using the fluoxetine device resulted in significantly higher absorption of fluoxetine into the wound tissues than bolus dosing using a pipette (p=0.0041, Fig. 2).

**Fig. 2.**
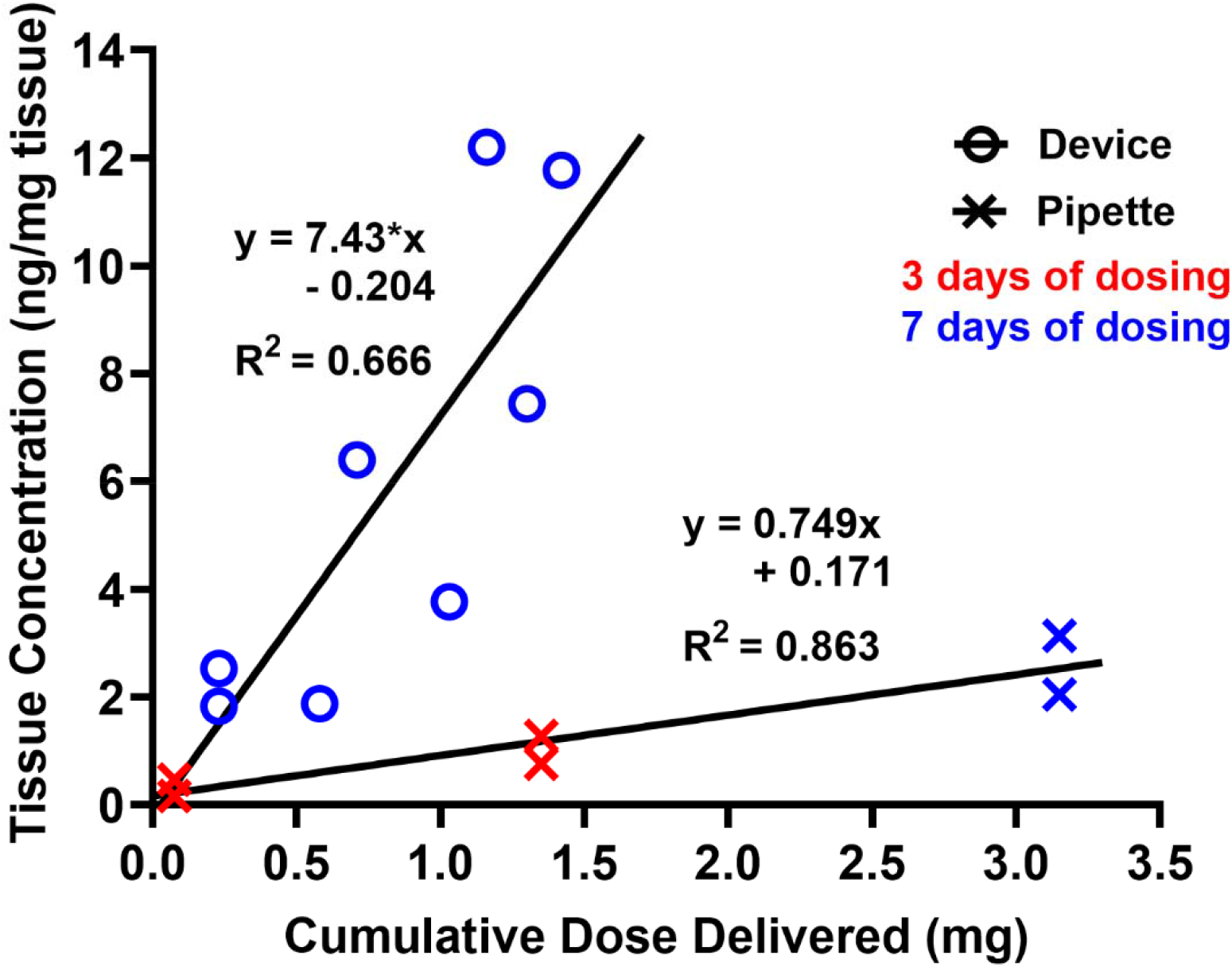
Tissue levels of fluoxetine in the wound following daily application using the experimental device or a pipette. Each data point represents one wound. Device, n=8 wounds; pipette, n=6 wounds.

### 3.2 Fluoxetine has a half-life of 0.99 days in the wound

After daily application of fluoxetine for 3 to 7 days using either the experimental device or a pipette (high- or low-dose), tissue was collected 1, 3, 4, 6, 8, or 16 days after the final application. The tissues were assayed for residual fluoxetine using HPLC, and the resulting concentrations are shown in Fig. 3a. The pharmacokinetic parameters of fluoxetine in the wound tissue were determined according to the equation

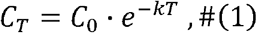

where *C*_*T*_ is the tissue concentration at time *T, C*_0_ is the tissue concentration following the final dose, and *k* is the elimination constant. Rearranging this equation yields

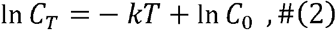

which is a linear equation. Accordingly, the natural logarithms of the tissue concentrations were plotted against time, and the elimination constant *k* and the tissue concentration *C*_0_ were determined graphically using the slope and y-intercept of the regression line[39] (Fig. 3b). The half-life *T*_*1*/*2*_ is the time required for one-half of the fluoxetine to be cleared from the wound tissue, such that

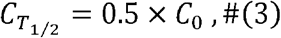

and substituting into equation (1) yields

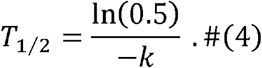

**Fig. 3.**
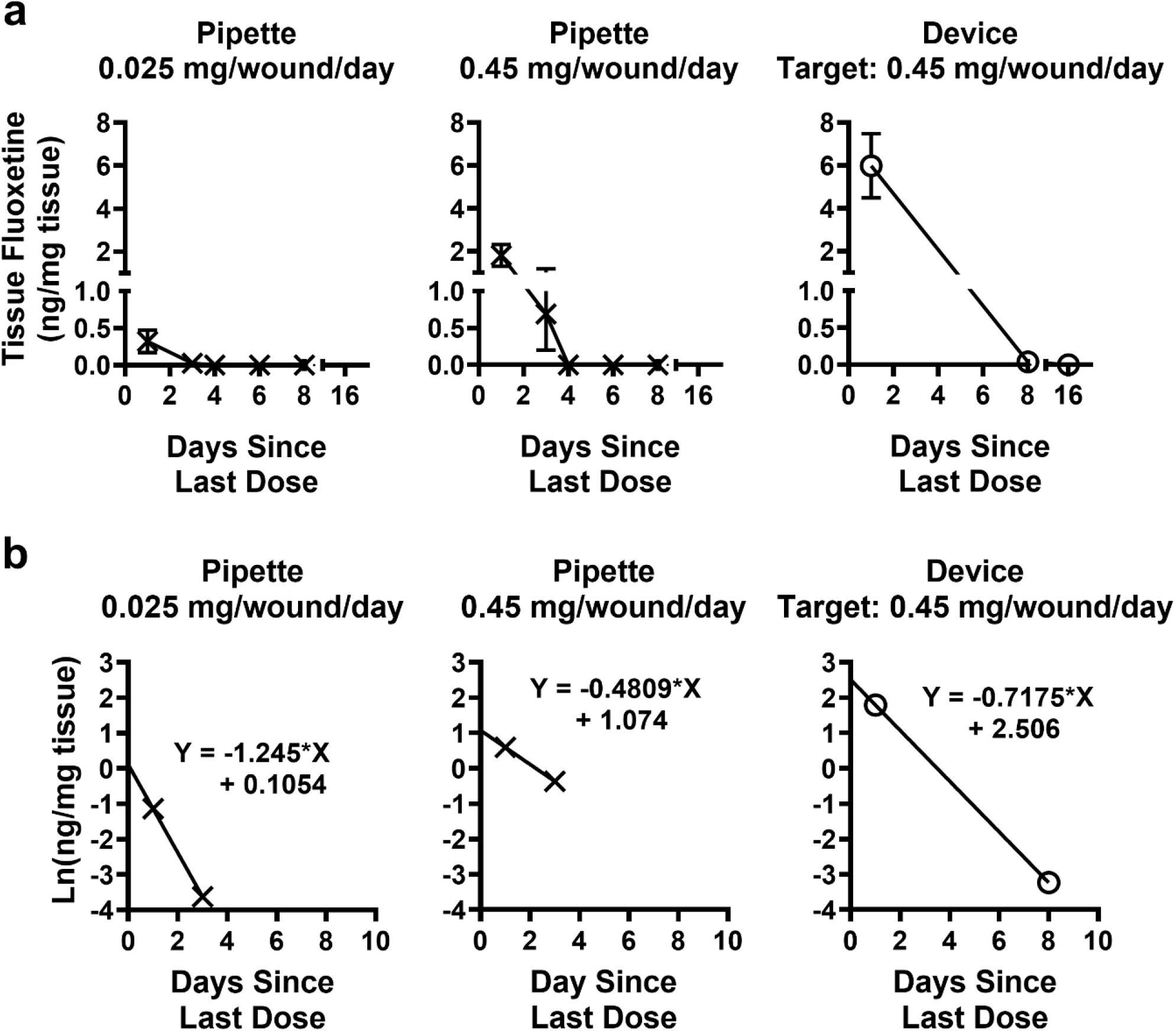
(a) Fluoxetine concentration in wound tissues collected on differing days post-treatment. Plots show mean ± standard error, n=2 to 8 wounds per point. (b) Plots of the natural logarithm of the tissue concentration across time for determination of the pharmacokinetic parameters.

The half-life of fluoxetine in the wound tissue was calculated for each dose regime and application modality, and the pharmacokinetic parameters are given in Table 1. The average half-life from the three conditions was 0.988 ± 0.256 days.

**Table 1.** Pharmacokinetic parameters of fluoxetine in the wound tissues.

|  | Elimination constant<br>(1/days) | Half-life<br>(days) | Concentration at T<sub>0</sub><br>(ng/mg tissue) |
| --- | --- | --- | --- |
| Low dose pipette | 1.245 | 0.5567 | 1.111 |
| High dose pipette | 0.4809 | 1.4413 | 2.926 |
| Fluoxetine device | 0.7175 | 0.9661 | 12.25 |

### 3.3 Topical fluoxetine is not absorbed into the systemic circulation

Fluoxetine was applied daily to two or four wounds, using either the experimental device or a pipette (high- or low-dose) for 3, 7, or 10 days, and plasma was collected one day after the final application. Plasma was analyzed for the presence fluoxetine and norfluoxetine, the primary active metabolite found in the blood of patients undergoing oral fluoxetine therapy. The limits of detection for our analysis were 7.4 ng/mL fluoxetine and 6.9 ng/mL norfluoxetine in pig plasma, and neither analyte was detected in the plasma after daily fluoxetine wound application (Fig. 4).

**Fig. 4.**
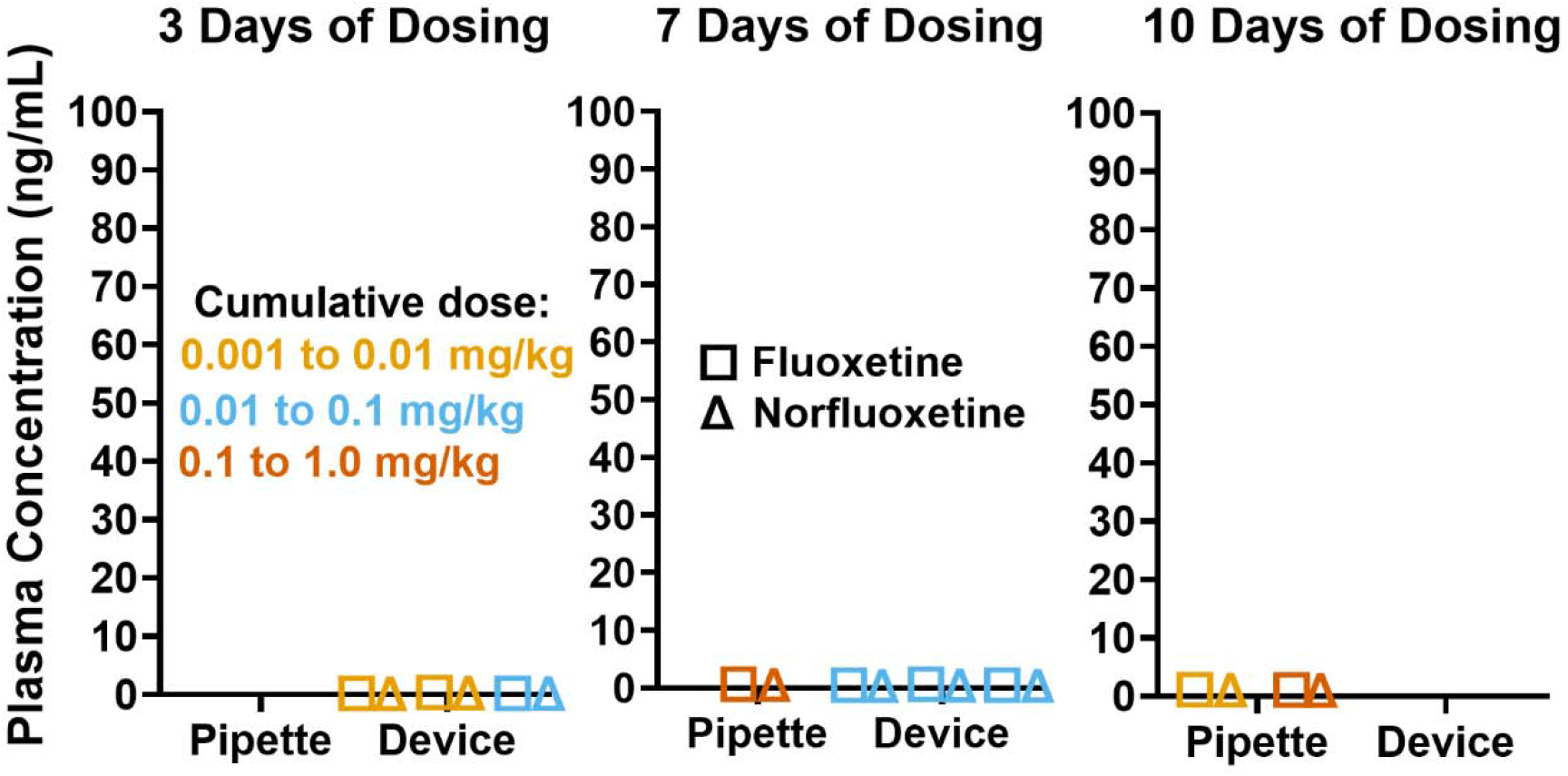
Fluoxetine and norfluoxetine concentrations in pig plasma (n=9 pigs) following topical fluoxetine wound treatment using the experimental device or a pipette. All results were below the limit of detection for the assay.

### 3.4 Topical fluoxetine does not alter systemic serotonin

Fluoxetine was applied daily to two or four wounds using either the experimental device or a pipette (high- or low-dose) for 3, 7, or 10 days. The plasma serotonin concentration was measured prior to wounding and again one day after the final fluoxetine application, and the change in plasma serotonin, Δserotonin, was calculated. For each pig, Δserotonin was compared to the cumulative fluoxetine dose delivered to the animal to determine whether changes in plasma serotonin were related to the topical fluoxetine wound application (Fig. 5). ΔSerotonin was not correlated with the duration of fluoxetine application or the cumulative fluoxetine dose administered to the pigs (3 days of dosing, Spearman’s R=0.103, p=0.90; 7 days, R=0.200, p=0.917; 10 days, R=-1.00, p=0.33).

**Fig. 5.**
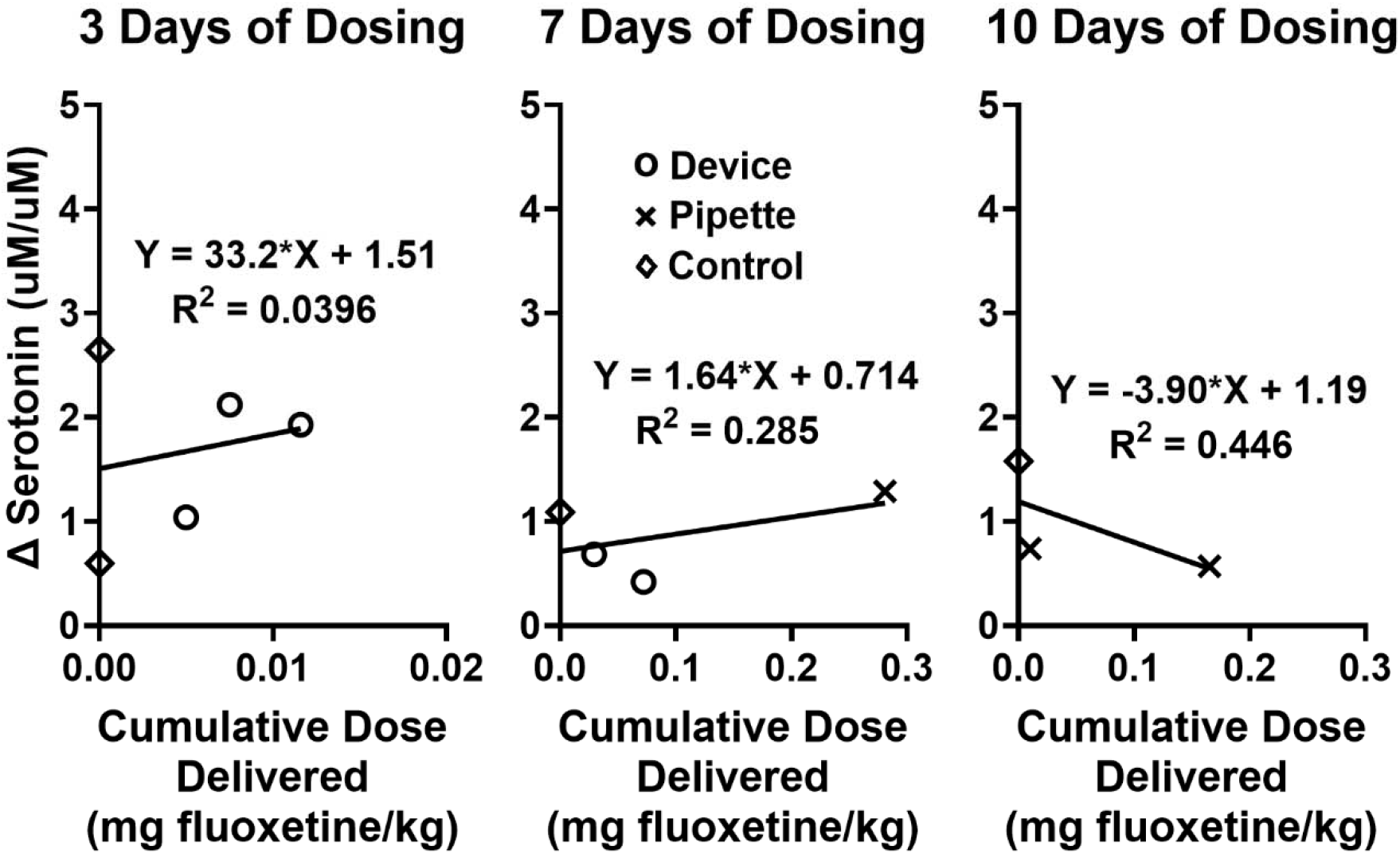
Changes in the plasma serotonin concentration after topical fluoxetine wound treatment using the experimental device or a pipette, plotted against the cumulative fluoxetine dose. Dose was normalized to the body weight of each pig. Each data point represents one pig. 3 days of dosing, n=5; 7 days, n=4; 10 days, n=3.

### 3.5 Topical fluoxetine increases the serotonin concentration in the wound

Topical fluoxetine was applied daily for 7 days using the experimental device or a pipette, and wound tissue serotonin was measured one day after the final application. Tissue serotonin and fluoxetine levels were compared, and serotonin was found to be increased in the presence of intermediate, but not high, tissue concentrations of fluoxetine (Fig. 6a and b).

**Fig. 6.**
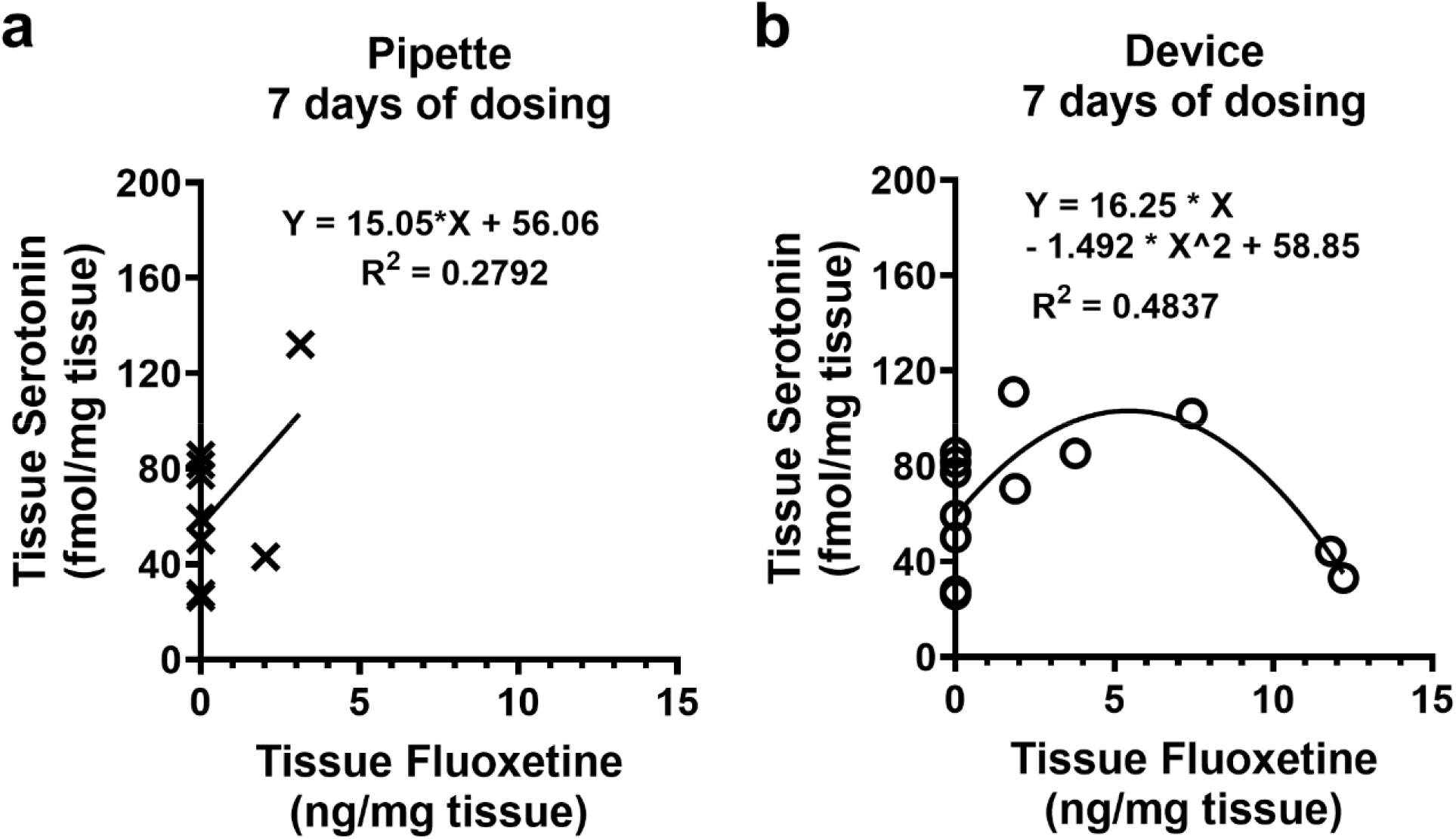
Wound tissue serotonin concentration after pipette (a) and device (b) topical fluoxetine wound treatment, plotted against the tissue fluoxetine level. Each data point represents one wound. Pipette, n=9 wounds; device, n=13 wounds.

## 4. Discussion

Interest in fluoxetine as a repurposed therapeutic to expedite wound healing is growing. Pre-clinical studies have demonstrated multiple pro-reparative functions: enhanced keratinocyte migration and neovascularization, reduced inflammation, and a shift in macrophage polarization towards a pro-reparative phenotype[15–19]. More recently, fluoxetine has been investigated for its antibacterial properties[9,10]. Fluoxetine exhibits bactericidal activity against multidrug-resistant (MDR) clinical isolates of *A. baumannii, E. cloacae, E. coli, E. faecalis, E. faecium, K. pneumoniae*, and *S. aureus*, with minimum inhibitory concentrations (MICs) ranging from 32 to 512 ug/mL; furthermore, the addition of fluoxetine at 1 ug/mL dramatically reduced the MICs of ciprofloxacin against all the clinical isolates tested[12]. Fluoxetine was also found to be bactericidal against MDR clinical isolates of *E. coli* and *P. aeruginosa*, with MICs of 102 and 161 ug/mL, respectively; in addition, the MICs of erythromycin against the same clinical isolates were significantly reduced in the presence of fluoxetine at 1/8^th^ of its MIC[13]. Most recently, fluoxetine was found to be bactericidal against several species of gram positive and gram negative bacteria, with MICs of 64-128 ug/mL and 8-512 ug/mL, respectively, and that fluoxetine combined with polymyxin B produced synergistic killing against 80% of the gram-negative strains tested[14]. Here, we demonstrate that metered delivery of topical fluoxetine using the experimental device results in significantly increased tissue concentrations of fluoxetine compared to bolus dosing using a pipette. Crucially, the experimental device produced a maximum concentration of 12.25 ug fluoxetine/mL tissue, above the MIC for one strain of *A. baumanii* and sufficient to produce synergistic killing with antibiotics against several MDR strains of gram-positive and gram-negative bacteria. Furthermore, the tissue concentration of fluoxetine was strongly correlated to the cumulative dose administered using the experimental device, which should allow physicians to reliably produce therapeutic concentrations in the wound tissue.

Future iterations of the device have the potential for closed-loop, wound stage-specific treatments and could identify the four classical stages: hemostasis, inflammation, proliferation, and maturation[40,41]. Each stage entails a distinct set of cellular and biochemical processes, and traditional standardized wound care protocols may not adapt to the evolving state of the wound. Integrated closed-loop systems have been developed to initiate treatment in real-time in response to wound assessments[28]; however, so far, no closed-loop devices exist to deliver adaptive treatments based on the wound stage[28]. To address this need, we developed a closed-loop system which supports AI-driven monitoring and decision making. The integrated camera within the device captures high-resolution images of the wound at regular intervals and transmits them to a central server, where they are processed by two models: a stage-evaluation AI model and a decision-making deep reinforcement leaning model (DRL). The models evaluate the wound stage and create a revised treatment plan which is transmitted to the device controller and implemented in real-time. More information about the closed loop functionality of the device is described in our other work[33].

The pharmacokinetics of fluoxetine as an oral therapy for psychiatric conditions have been well studied since the 1970’s[42]. Fluoxetine is lipophilic and has a large volume of distribution (V_d_), indicating extensive accumulation in tissues[43–45]. For this reason, fluoxetine has a long elimination half-life of 1-4 days[44,46]. However, the pharmacokinetics of fluoxetine in wound skin are not well understood. Transdermal delivery systems have been developed[47– 49] but, due to the barrier function of the intact epidermis, these studies provide limited insight into the pharmacokinetics of fluoxetine in wounds where the epidermis is not present and delivered drug has access to the dermal vasculature. In addition, although some groups used carriers to increase tissue absorption, skin fluoxetine levels were not measured. We found that fluoxetine had a half-life of 0.988 ± 0.256 days in the wound tissue, consistent with the elimination half-life measured in the blood after oral dosing[44,46]. Determination of the half-life of fluoxetine in wound tissue may inform the frequency of topical application. Once-daily oral dosing with fluoxetine is often used for the treatment of psychiatric conditions[50], and this also may be suitable for fluoxetine topical wound therapy. In multiple-dosing, a fixed dose is applied repeatedly at intervals of one half-life; typically, steady state concentrations occur after 5 doses and are twice the concentration achieved after the initial dose[39]. If fluoxetine is applied too infrequently a therapeutic concentration may not be achieved, while accumulation to higher than desirable levels can occur if application is too frequent[39]. Thus, the advantage of the experimental device described here is that its programmable dosing can be adjusted to deliver fluoxetine at precise intervals, offering an advantage over manual dosing where accurate timing of each dose may be a challenge. In addition, automated dosing from the device may simplify wound treatment and improve patient adherence to the treatment regimen.

Fluoxetine is considered to have a good safety profile[51,52], exhibiting high specificity for serotonin uptake sites and low affinity for serotonin receptors and the uptake sites of norepinephrine[53] and dopamine[54]. Common side effects are limited and life threatening side effects such as cardiotoxicity and CNS toxicity are rare due to the lack of monoamine receptor antagonism[44]. In addition, drug-drug interactions are possible and have become important selection criteria when considering fluoxetine use[55,56]. Fluoxetine inhibits the isoenzymes CYP2D6[57–59] and, to a lesser extent, CYP3A3/4[60] and produces impairments in the clearance of drugs metabolized by these cytochromes, such as TCAs, neuroleptics, alprazolam, and carbamazepine[44]. Due to the possibility of side effects and drug-drug interactions, we evaluated systemic absorption of fluoxetine after topical administration to the wound bed, by analyzing plasma levels of fluoxetine and its metabolite norfluoxetine after application to porcine skin wounds. Fluoxetine and norfluoxetine were not detectable in the plasma after 3, 7, or 10 days of daily topical wound application. The therapeutic window for the combined concentration of fluoxetine and norfluoxetine in the blood of patients undergoing oral fluoxetine therapy is 120 to 500 ng/mL[61], and the results of the present study suggest that side effects and drug-drug interactions are unlikely to result from topical application of fluoxetine to wounds.

When fluoxetine is used as a topical therapy for wounds, effects should ideally occur only in the wound site. Fluoxetine has the potential to alter systemic serotonin levels. Plasma serotonin increases (∼60%) shortly after oral fluoxetine administration (7 to 10 hours)[62], but then decreases after long term (8 weeks) daily treatment[63]. To investigate the possibility of unwanted systemic effects in the experimental animals, plasma serotonin was measured prior to each experiment and again at the experimental endpoint. We found that changes in plasma serotonin levels were not correlated with the cumulative doses of fluoxetine delivered to the animals on their wounds. Taken together with the lack of detectable fluoxetine and norfluoxetine in the plasma, these results indicate that topical fluoxetine wound treatment is unlikely to produce systemic effects.

Fluoxetine is thought to improve skin wound healing through the modulation of serotonin signaling. The primary source of serotonin in the skin is in platelets, but serotonin is also synthesized and stored in skin cells[64–66]. Serotonergic neurons are not present in the skin[64]. Fluoxetine prevents cellular reuptake by blocking the serotonin transporter SERT, which is expressed on immune cells[65], keratinocytes[67,68], fibroblast[69], and Merkel cells[66]; SERT is also expressed on platelets, and fluoxetine has been shown to reduce the platelet-bound fraction of serotonin in the blood[70,71]. It follows that fluoxetine may increase extracellular serotonin in the wound skin. Serotonin receptors comprise a family of G protein-coupled and ligand-gated ion channel receptors expressed on cell surfaces[72], and an increase in unsequestered serotonin may enhance signaling and produce beneficial effects on healing. In in-vivo scratch wound models using primary human keratinocytes, serotonin accelerated closure in a dose dependent manner[16,69], and other groups have found that serotonin skewed human macrophage polarization towards a less inflammatory phenotype[73] and reduced the expression of the pro-inflammatory cytokines TNFa and IL6[74]. Fluoxetine treatment produced similar effects in an excisional wound model using diabetic mice, enhancing re-epithelialization, reducing the expression of TNFa and IL6, and shifting macrophage polarization towards a less inflammatory and more pro-reparative phenotype[16]. Here, we report for the first time the serotonin modulating effects of fluoxetine in the skin, where fluoxetine at intermediate, but not high, tissue concentrations increased the total tissue level of serotonin. This suggests that in the case of non-infected wounds, lower tissue concentrations of fluoxetine may be more beneficial for healing. However, more work is needed to investigate changes in the proportions of free and sequestered serotonin in the wound tissue independent of changes in the total amount. A key advantage of the experimental device is programmable dosing which can be adjusted depending on the specific use case; higher doses can be applied to elicit an antimicrobial effect in the case of infected wounds, and lower doses can be delivered to stimulate healing by modulation of tissue serotonin in the case of uninfected wounds.

## 5. Conclusion

In this study, we demonstrated that a bioelectronic wound treatment device can effectively deliver topical fluoxetine to wounds with minimal risk of off-target effects. The experimental device produces higher tissue concentrations of fluoxetine at lower cumulative doses compared to bolus dosing, which may lead to greater efficacy when fluoxetine is used as an adjunctive therapy for infected wounds. We established basic pharmacokinetic parameters of fluoxetine in the wound tissue and investigated the effects of fluoxetine on the endogenous serotonin system in the skin. The device may simplify wound treatment by reducing the burden for daily drug application, possibly increasing adherence to a prescribed treatment regimen. Recent iterations of the device incorporate closed-loop feedback control using integrated imaging and machine learning to produce real-time adaptive treatments based on the predicted wound stage, and work is ongoing to optimize the design for commercialization and clinical use.

## CRediT authorship contribution statement

**Anthony Gallegos:** Writing – original draft; writing – review and editing; conceptualization; investigation; methodology; formal analysis. **Houpu Li:** Writing – review and editing; investigation; methodology; software. **Hsin-ya Yang:** Writing – review and editing; investigation; methodology. **Guillermo Villa-Martinez:** Investigation; methodology. **Itidal Bazzi:** Investigation. **Samhitha Sathyanarayanan:** Investigation. **Narges Asefifeyzabadi:** Investigation; software. **Prabhat Baniya:** Methodology; software. **Wan Shen Hee:** Investigation; software. **Mo Siadat:** Investigation. **Elizabeth Chang:** Investigation. **Sriansh Pasumarthi:** Investigation. **Elham Aslankoohi:** Project administration. **Mircea Teodorescu:** Conceptualization; methodology; resources; funding acquisition. **Marcella Gomez:** Conceptualization; software; methodology; funding acquisition. **Marco Rolandi:** Supervision; conceptualization; methodology; resources; funding acquisition; writing – review and editing. **Rivkah Isseroff:** Supervision; conceptualization; methodology; resources; funding acquisition; writing – review and editing.

## Ethics approval statement

Animal studies were performed according to the guidelines of the Association for Assessment and Accreditation of Laboratory Animal Care (AAALAC) and the NIH Guide for the Care and Use of Laboratory Animals, and were approved by UC Davis Campus Veterinary Services and the UC Davis Institutional Animal Care and Use Committee (IACUC protocol # 23353).

## Acknowledgements

This research is sponsored by the Defense Advanced Research Projects Agency (DARPA) and the Advanced Research Projects Agency for Health (ARPA-H) through Cooperative Agreement D20AC00003 awarded by the US Department of the Interior (DOI), Interior Business Center. The content of the information does not necessarily reflect the position or the policy of the government, and no official endorsement should be inferred.

We thank Justine Arizabal for creating the wound illustrations used in this paper.

## Data availability

Data supporting this study are available from the corresponding author upon reasonable request.

